# The influence of muscle mass on the coordination required for efficient movement

**DOI:** 10.64898/2026.04.30.722018

**Authors:** Ameur Latreche, Stephanie A. Ross, Taylor J.M. Dick, Nicolai Konow, Andrew A. Biewener, James M. Wakeling

**Affiliations:** Department of Biomedical Physiology and Kinesiology, Simon Fraser University, Canada; Faculty of Kinesiology, University of Calgary, Canada; School of Biomedical Sciences, The University of Queensland, Australia; Department of Biological Sciences University of Massachusetts Lowell, MA, USA; Organismic and Evolutionary Biology Harvard University Cambridge, MA, USA

**Keywords:** muscle inertia, mass gravity, Hill-type muscle model, musculoskeletal simulation, muscle coordination

## Abstract

Muscle efficiency decreases with increasing size, largely due to a relative decrease in its mechanical output. Muscle mechanical output depends on its activation, strain, and strain rate and thus varies between different muscles within a limb during locomotion. Distinct muscle coordination patterns are required for efficient cycling, and so we would expect that the coordination patterns for efficient cycling or indeed locomotion would change across animal sizes. We tested whether muscle coordination would change with muscle size using data derived from human cycling: this paradigm allowed for controlled changes in both crank torque and cadence, allowing the multifactorial problem of muscle power output to be decomposed. We used kinematic and pedal data from 12 cyclists undergoing steady pedalling at cadences from 80 to 140 r.p.m. and generated musculoskeletal simulations of their movements. We introduced novel multisegment muscle models in the simulation that incorporated the internal muscle mass and thus accounted for the scaling effects of muscle tissue inertia. We solved the simulations for the muscle activity that was required to minimise the metabolic cost during cycling for each condition. The masses of the muscle models were scaled across five orders of magnitude. The predicted muscle activations were classified by Principal Component analysis to identify whether the coordination of muscle activity was modulated across models with different sized muscles. Analysis of variance revealed significant changes in coordination at the large-scale factors. This study shows how the coordination of muscle activity during locomotion will likely change across a range of body sizes due to the non-linear effects of the inertial mass within the muscle tissues.

## Introduction

Muscle-driven movement emerges from the interaction between neural commands, musculotendon mechanics, and task dynamics. Terrestrial mammals display an extraordinary range of body sizes, from mice to elephants. Yet all must generate and control locomotion despite vastly different mechanical demands [34, 35]. Because muscle mass scales strongly with size [36, 83, 84], larger mammals possess muscles with substantially greater inertial and gravitational loads, which can alter the dynamics of shortening and lengthening by requiring additional force to accelerate internal tissue and by modifying effective force transmission during rapid, cyclic movements [36]. However, the force a muscle can produce does not scale as much as mass as animals increase in size. At the same time, it is unclear to what extent larger animals mitigate these demands through musculoskeletal design features [80] and control strategies (e.g., tendon compliance [85], muscle architecture, or activation timing), or whether increased muscle inertia remains an unmodeled contributor to coordination.

The inertia of the muscle tissue, or more generally the soft-tissues of the lower extremities results in soft-tissue motion that is not directly coupled to the skeletal movement in the limb segments. This “wobbling mass” phenomenon affects the force-transients at the joints [14] and the ground reaction force [56] during walking and running. It has been proposed that the muscle activation may be “tuned” to minimize soft-tissue resonance that occurs after the foot contact with the ground [55], this can be achieved through shifting the natural frequency or damping of the soft-tissues [65], with energy dissipation occurring at the cross-bridges [16]. Recent studies have directly tracked soft-tissue vibrations within the muscles of humans [54], and carefully recreated the impact phenomena and characterised the vibration modes in the gastrocnemii of the rat ex vivo [15, 16]. However, the effect of wobbling mass on mechanical output from muscles has not been explicitly tested, and the presence of intramuscular mass with its effects on inertia have rarely been considered in the muscle models used in the musculoskeletal simulations for the study movement and locomotion [63].

Classical Hill-type muscle models capture key static and velocity-dependent properties of muscle–tendon units (MTUs), but are typically *massless*: forces transmit quasi-statically through series elastic elements without inertial states along the muscle–tendon pathway [1, 2]. This simplification underpins much of contemporary musculoskeletal simulation and optimization, yet it ignores the effect of intramuscular mass on rates of force development [66], force, work [68, 69] and efficiency [67] of muscle contractions, and can obscure how accelerations and gravitational loading along the muscle–tendon pathway shape coordination, particularly in cyclic tasks where fibre and tendon velocities can be substantial.

Musculoskeletal models simulations provide a principled way to *predict* activation strategies for a given movement by solving muscle redundancy (e.g., via static optimization) or by performing predictive/optimal-control simulations, thereby linking task objectives and whole-body mechanics to muscle-level activation patterns [37]. This approach is increasingly used in comparative locomotion: for example, simulations have been used to estimate functional groupings and activation patterns in mouse locomotion [38], to predict muscle excitations and work in ostrich walking and running [39], and to build and analyze musculoskeletal simulations for extinct taxa to infer plausible locomotor mechanics in dinosaurs and other fossil vertebrates [40]. Related scaling-focused studies have also examined how predicted movement performance and coordination change as models are scaled across large ranges of body mass, typically by modifying body/segment geometry and inertial properties [41]. In parallel, most studies probing “mass effects” on coordination varied external loads or segment inertias and infer changes from EMG or joint-level dynamics [5, 6, 17, 18, 19, 20, 21]. However, in all these frameworks the MTUs are almost always represented with *massless* Hill-type actuators, meaning that inertial and gravitational demands *within* the muscle–tendon pathway are not explicitly represented [1, 2]. Consequently, it remains unclear how explicitly modelling musculotendon inertia would alter predicted activation patterns—and, downstream, whether coordination modes primarily reflect amplitude scaling of a dominant activation pattern, timing/redistribution among synergists, or both, when *musculotendon inertia is explicitly modeled*.

Most musculoskeletal simulation studies use multibody models with massless Hill-type muscles [1, 19, 24, 42, 52, 62]. Although these studies provide new insights into many biomechanical and clinical questions, the classic 1D Hill-type muscle model approach has several drawbacks (due to its simplifications, which do not account for distributed tissue mass). Furthermore, 3D muscle deformations during contractions, combined with complex changes in internal muscle architecture and 3D aponeurotic deformation, cannot be represented [59, 60, 61]. Muscles must work against their own inertia during contractions, which affects muscle performance [57] an aspect that is not taken into account in classic Hill-type muscle models [58, 62]. The use of 1D mass-enhanced muscle models is a step in the right direction, as it allows the influence of muscle inertia to be investigated under defined boundary conditions. So, Chen et al. [22] examined how muscle mass influences the muscle’s own contractile properties during various movement tasks, with muscle masses scaled over six-orders of magnitude. However, they focused on abstracted muscles rather than simulating the entire human lower limb musculoskeletal system. Here, we consider how the complex coordination of the activities between muscles changes with muscle size while the model strives to achieve locomotor patterns with a low metabolic cost. Understanding how scale affects biomechanical results in musculoskeletal models will allow more accurate models to be developed [27, 28, 29, 30, 31, 45].

*Muscle synergies* are groups of muscles that share common activation patterns over space or time [3, 4, 10]. In cyclic lower-limb tasks (e.g., pedaling-like movements), synergy analyses often identify a dominant coordination pattern that can be further modulated across muscles over the cycle [12, 64]. While coordination patterns are known to depend on cadence and mechanical context [11, 12, 25], it remains unclear whether and how these patterns reflect *inertial properties of the musculotendon system itself*, as opposed to only segment-level dynamics or external torques.

The goal of this study is to implement *muscle mass* within a whole-body musculoskeletal model and test whether explicitly including inertia in Hill-type muscle–tendon dynamics is *functionally important* for predicted coordination patterns. To do so, we discretized each MTU into a one-dimensional chain of lumped masses and elastic elements (preserving internal force transmission, tendon stretch, and pennation geometry) and embedded this mass-augmented MTU within a muscle redundancy solver (MRS) framework [8]. Unlike the canonical massless MTU, segment positions and velocities now evolve under Newtonian dynamics (including gravitational terms), so contractile forces must accelerate distributed muscle–tendon mass rather than being transmitted in a quasi-steady manner. We focused on human cycling because human musculoskeletal models and cycling simulations are more mature for walking (e.g., EMG, kinematics/kinetics, and muscle–tendon/fascicle dynamics), providing a stronger basis for validation than is typically available across a broad range of species [46, 47]. Cycling is also an ideal testbed because cadence and crank power/torque can be manipulated systematically, enabling a wide range of fibre shortening/lengthening velocities and force demands [48, 49]. The coordination patterns depend on the overall cycling efficiency [70], paralleling our approach to minimize the metabolic cost of cycling. Finally, cycling elicits rhythmic, cyclic patterns of muscle activation that share modular features with other locomotor behaviors, supporting translation of mechanistic findings beyond this specific task [50]. Accordingly, we systematically vary the *muscle mass parameter* and *cadence*, solve the MRS across subjects and conditions, and quantify the resulting coordination as changes to the predicted time-varying muscle activations.

We tested two hypotheses about how the muscle mass influences the muscle activations and coordination between muscles.

*H1 (Increasing muscle mass will increase the muscle activations)*. We hypothesized that a larger muscle mass-scale would result in a general increase in muscle activations for a given movement, as the muscles have to do greater internal work to overcome their relatively greater inertia.

*H2 (Increasing muscle mass will modulate the muscle coordination)*. We hypothesized that a larger muscle mass-scale would result in muscle-specific changes in muscle activation, because each muscle has a unique size, velocity and acceleration. This change in muscle-specific activation would be observed by a change in the time-varying balance of activations that would modulate the general muscle coordination pattern.

## Methods

### 2.1 Approach to the problem

The purpose of this study is to examine the effect of muscle mass on the predicted activations and muscle coordination patterns during movement. Here we have taken a modelling approach: the simulations track experimental data collected from human cycling, and we implement the ability to account for (and adjust) the mass within each muscle. Note that in this context the simulations are all human-sized (with the muscle MTU lengths, velocities and the required joint moments all being human-sized), however, the muscles within the models have masses that are scaled over a range of five orders of magnitude.

### 2.2 Experimental measures of cycling

We used previously published cycling data [23] from 12 healthy people (6 men, 6 women: aged 24-43 years, height 1.67-1.83 m, mass 58.0-82.8 kg): detailed data in Table S1. The institutional Ethics Review Boards at Simon Fraser University approved the study protocols, and the participants provided informed consent. We used data from four different cycling conditions: 3 conditions were for cycling against a crank torque of 12 N m, and at cadences of 80, 120 and 140 r.p.m. (referred to as 80T12, 120 and 140, respectively), and one condition was at a cadence of 80 r.p.m. and crank torque of 40 N m, referred to as 80T40. Data were analysed for three non-sequential pedal cycles within each trial, and each of these pedal cycles started and finished with the right crank at top-dead-centre and had its time expressed as a percentage of that cycle.

### 2.3 Musculoskeletal Model

A musculoskeletal model (21 degrees of freedom (DoF): 6 at the pelvis, 7 in each leg, 0 in the torso and upper body, and an extra coordinate related to the cycling/crank mechanism) was scaled to the dimensions of each participant, and the kinematic and crank load data from the cycling trials were used to run musculoskeletal model simulations [24]. Each model was driven by 43 muscle actuators that generate moments about the lower limb joints. The default muscle actuators (Hill-type muscle models) were replaced by mass-enhanced muscle Hill-type muscle models that were formulated in the following manner.

Conventional Hill-type muscle models typically neglect the explicit inertial effects of muscle tissue, treating the contractile element as massless and assuming instantaneous transmission of force to the tendon. While this assumption simplifies the dynamics, it overlooks the distributed mass of the muscle belly, which can influence transient force responses, energy transfer, and the stability of musculoskeletal simulations, especially when muscle size is altered [51, 66]. In the present formulation, we explicitly represented the inertial properties of the muscle belly (following [66, 69, 22]) by discretizing each muscle belly into *n*_seg_ serial lumped-mass elements. Each element has a contractile segment (based on a Hill-type muscle model) in series with a point-mass *m*_seg_

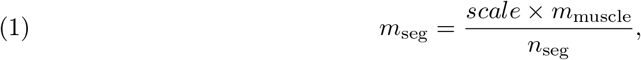

where *m*_muscle_ is the muscle-specific mass obtained from subject-specific OpenSim scaling and *scale* is the mass-scale factor. The state of each point-mass is defined by its one-dimensional position *p*_*i*_(*t*) along the muscle-tendon unit MTU line of action and its velocity *v*_*i*_(*t*).

In this study, we used *n*_seg_ = 5 segments per MTU to balance model fidelity and computational tractability. In pilot tests with a larger number of segments, the predicted state and activation trajectories changed negligibly for the tasks considered, whereas the computational cost increased substantially. For each segment *s* of muscle *m*, the translational dynamics were described using Newton’s second law along a one-dimensional internal muscle axis:

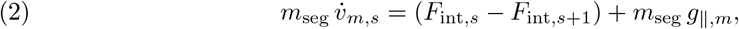

where *m*_seg_ is the segment mass, *F*_int,*s*_ and *F*_int,*s*+1_ are the internal forces at the proximal and distal interfaces of the segment, and *g*_*∥,m*_ is the component of the gravity vector projected onto the instantaneous MTU line of action:

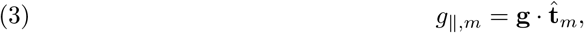

with **g** the gravity vector expressed in the global frame and 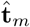 the unit vector defining the instantaneous muscle direction. Thus, gravity was not applied as a full axial load, but only through its projection along the internal one-dimensional muscle coordinate.

The internal forces at the boundaries of the MTU were provided by the Hill-type muscle– tendon formulation. At the proximal boundary, the lumped system received the muscle-side force along the line of action,

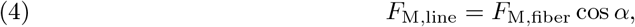

where *F*_M,fiber_ is the fiber force and *α* is the pennation angle. At the distal boundary, the lumped system was connected to the tendon force *F*_T_. Internal interface forces between adjacent segments were introduced as optimization variables, allowing force transmission through the discretized muscle belly.

State updates were implemented using an explicit Euler scheme:

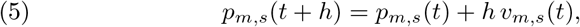

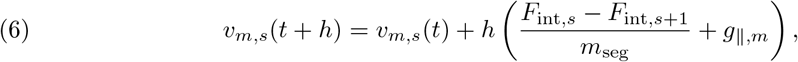

with mesh step *h*. In the numerical implementation, this formulation ensures that changes in activation, tendon force, pennation, and projected gravity propagate through the muscle belly via physically consistent internal force imbalances.

The force–length and force–velocity relationships of the contractile element, as well as the series-elastic and parallel-elastic components of the Hill-type actuator, followed the default muscle–tendon relations of the Hill-type muscle model formulation used in the original MuscleRedundancySolver framework [8]. The maximum unloaded contraction velocity was set to 10 *l*_*M*0_ s^*−*1^.

### 2.4 Predictions of muscle activity and coordination

Musculoskeletal simulations were run for *N* = 695 pedal cycles from 720 (12 participants x 4 conditions x 3 pedal cycles for each trial x 5 mass scale factors). The inverse kinematics and inverse dynamics were used in a muscle redundancy solver to predict the activations of the lower limb muscles [8]. The nominal human-sized muscles were considered as mass-scale 1, we additionally tested muscle mass-scales at 0.01, 25, 70, and 125. These scales span the approximate range of mammalian masses from a shrew (scale 0.01 [78]) to the extinct giant hornless rhinoceros (scale 125 [79]). By keeping the inertial dynamics one-dimensional along the line-of-action of each muscle within a CasADi/IPOPT framework (CasADi is an open-source symbolic/automatic-differentiation framework used to formulate nonlinear optimization problems, which can then be solved by IPOPT, an open-source interior-point solver for large-scale nonlinear programming [32, 33]), the method remains computationally tractable while capturing the dominant inertial effects along the MTU line of action.

#### Cost function

The optimal control problem was formulated to minimize a weighted combination of physiological effort, reserve-actuator effort, contraction-velocity regularization, metabolic energy expenditure, activation smoothing, and internal lump-velocity regularization over the discretized time mesh. In line with direct-collocation muscle-redundancy formulations, the objective combines muscle-control penalties commonly used in predictive and tracking simulations with additional regularization terms required by the present mass-enhanced extension [8, 53].

In discrete form, the objective function was written as

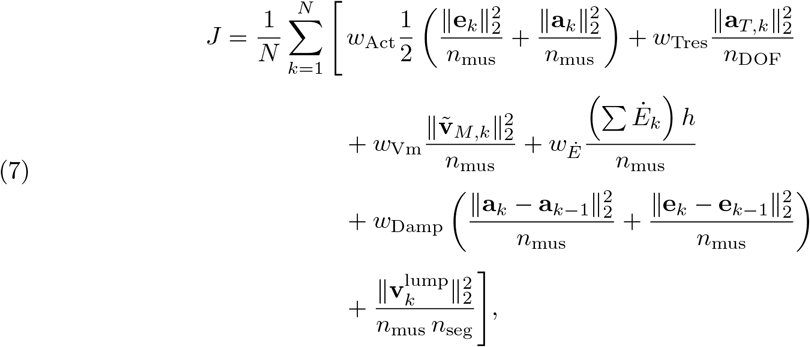

where **e**_*k*_ and **a**_*k*_ denote the excitation and activation vectors at mesh point *k*, **a**_*T,k*_ denotes the reserve-actuator controls, 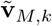 is the vector of normalized fibre contraction velocities, *Ė*_*k*_ is the vector of metabolic power rates, **v**^lump^ contains the internal lump velocities, *h* is the mesh step, *N* is the number of mesh intervals, *n*_mus_ is the number of modelled muscles, and *n*_seg_ is the number of lumped segments per muscle. An activation-smoothing term was applied from the second mesh interval onward.

The first term penalizes muscle effort through the squared magnitudes of excitations and activations. The second term penalizes reserve-actuator usage, thereby discouraging reliance on non-physiological auxiliary torques. The third term regularizes normalized fibre contraction velocity to avoid unrealistically large values. The fourth term minimizes metabolic energy expenditure using the Minetti–Alexander metabolic formulation [53]. The fifth term penalizes rapid changes in excitations and activations to promote temporally smooth control solutions. The final term regularizes the velocities of the internal lumped masses introduced in the one-dimensional muscle-inertia model, thereby improving numerical robustness while limiting unrealistically large internal motions. We settled the cost function (equation 2.7), with the weights shown in (2.8), to obtain a balance between computation efficiency and produced realistic cycling kinematics.

In the simulations reported here, the default weights were

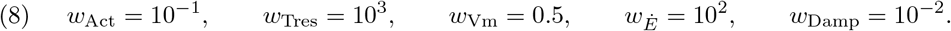

The lump-velocity regularization term was included with unit coefficient in the implemented objective. Under this weighting strategy, metabolic energy minimization remained the dominant physiological objective, whereas the remaining terms acted primarily as regularization terms to maintain feasible, smooth, and physiologically plausible muscle-force solutions in the presence of reserve actuation and internal muscle-mass dynamics [8].

### 2.5 Statistical analysis

Activations were calculated for 43 muscles for 100 interpolated time points for each pedal cycle (crank top-dead-centre to top-dead-centre), leading to 4300 activation values predicted across the set of 695 simulations. This large, multivariate data set was reduced to a series of coordination patterns using Principal Component (PC) analysis, following [64]. For each simulated pedal cycle there were 4300 calculated activations (43 muscles * 100 time points). These were arranged into a 695 by 4300 matrix. The mean value for each column (muscle and time-point) was subtracted, and the covariance matrix calculated from this detrended matrix. The eigenvectors from the covariance matrix form the PC weights and can be visualised as the principal coordination patterns. The loading scores for each PC describe the extent to which the PC weights contributed to each simulated pedal cycle [64].

The statistical effects of the pedalling condition and simulated muscle mass on the predicted muscle coordination patterns were determined using univariate linear mixed-effects analysis of variance for the loading scores of each PC. *Mass scale* (five levels: 0.01, 1, 25, 70, 125) and *Condition* (four levels: 80T12, 80T40, 120, 140 rpm) were treated as fixed factors, and *Participant* was included as a random factor. Where there was a significant effect (p<0.05) of mass-scale or condition on the PC loading score then the estimated marginal mean values for those conditions were calculated. If there was no significant effect then the global mean loading score was calculated for that factor.

## Results

The mean activation patterns for all participants for selected muscles for the four cycling conditions can be seen in Figure. 2. A total of 695 musculoskeletal simulations of cycling were evaluated. The mean of the predicted muscle activation patterns can be seen in Figure. 3 for ten select muscles. There was a general increase in the muscle activation as the cycling cadence increased. The mean activation pattern (Figure. 3A) was broadly similar in shape to the activation patterns from the separate cycling conditions (Figure. 2), and the PC analysis showed that the first five PCs explained 62.94% of the variance in activation pattern from this mean. The weights for PC1-5 are shown in Figure. 3B-F.

**Figure 1.**
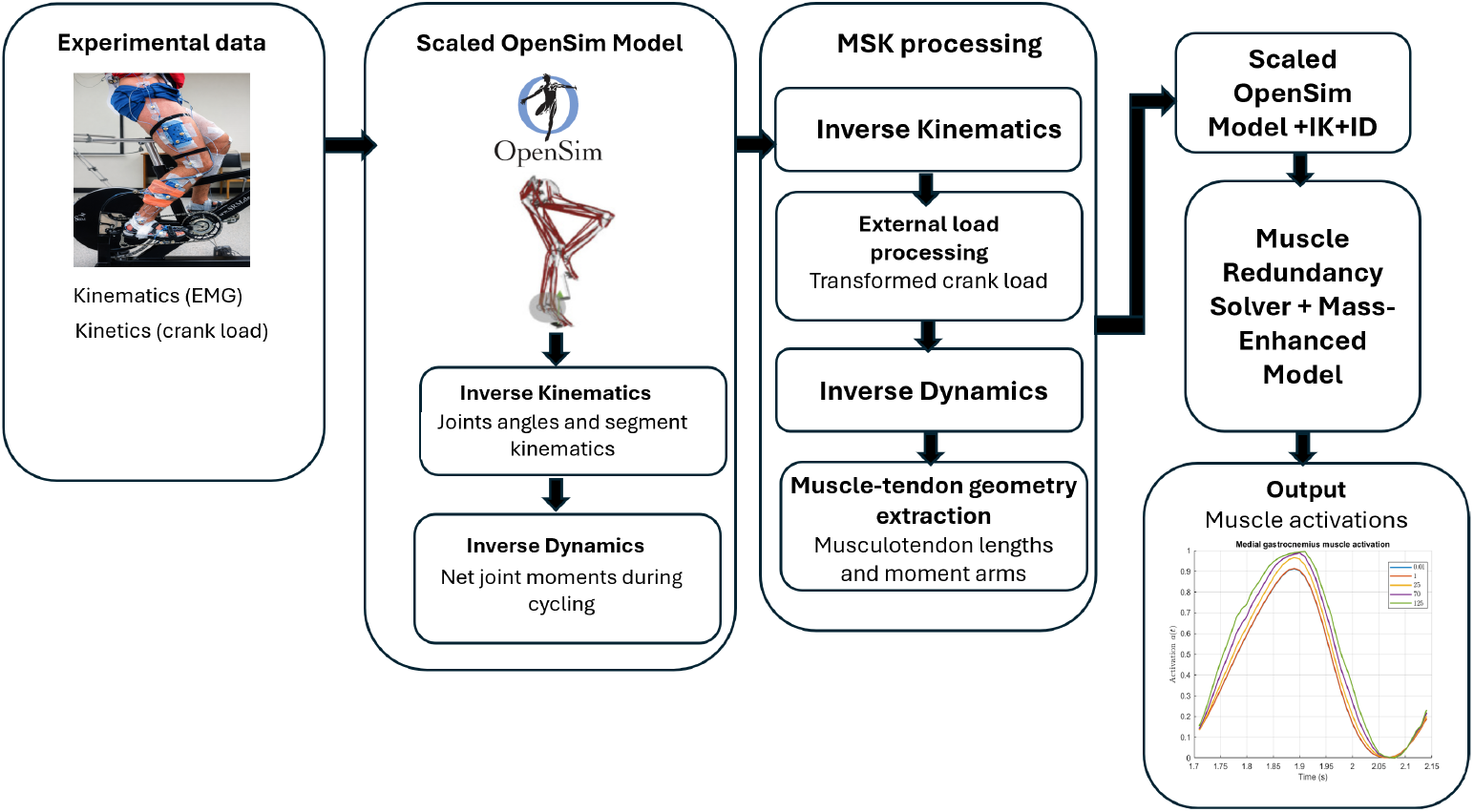
Workflow for muscle activation estimation during cycling a mass-enhanced muscle redundancy solver.

**Figure 2.**
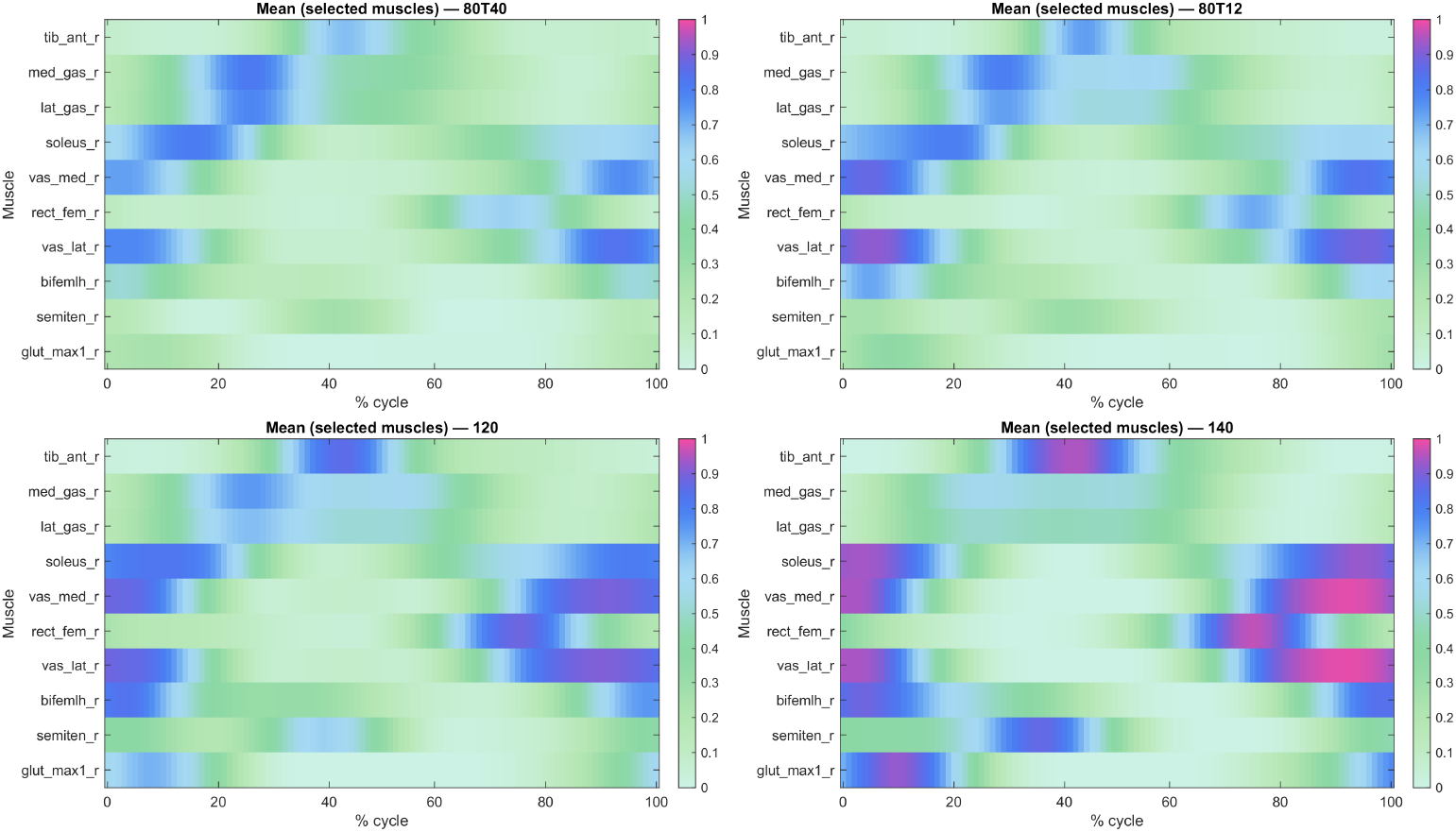
The means of predicted muscles activations for all participants for selected muscles for the four tested cycling conditions. 3 conditions were for cycling against a crank torque of 12 N m, and at cadences of 80, 120 and 140 r.p.m. (referred to as 80T12, 120 and 140, respectively), and one condition was at a cadence of 80 r.p.m. and crank torque of 40 N m, referred to as 80T40

**Figure 3.**
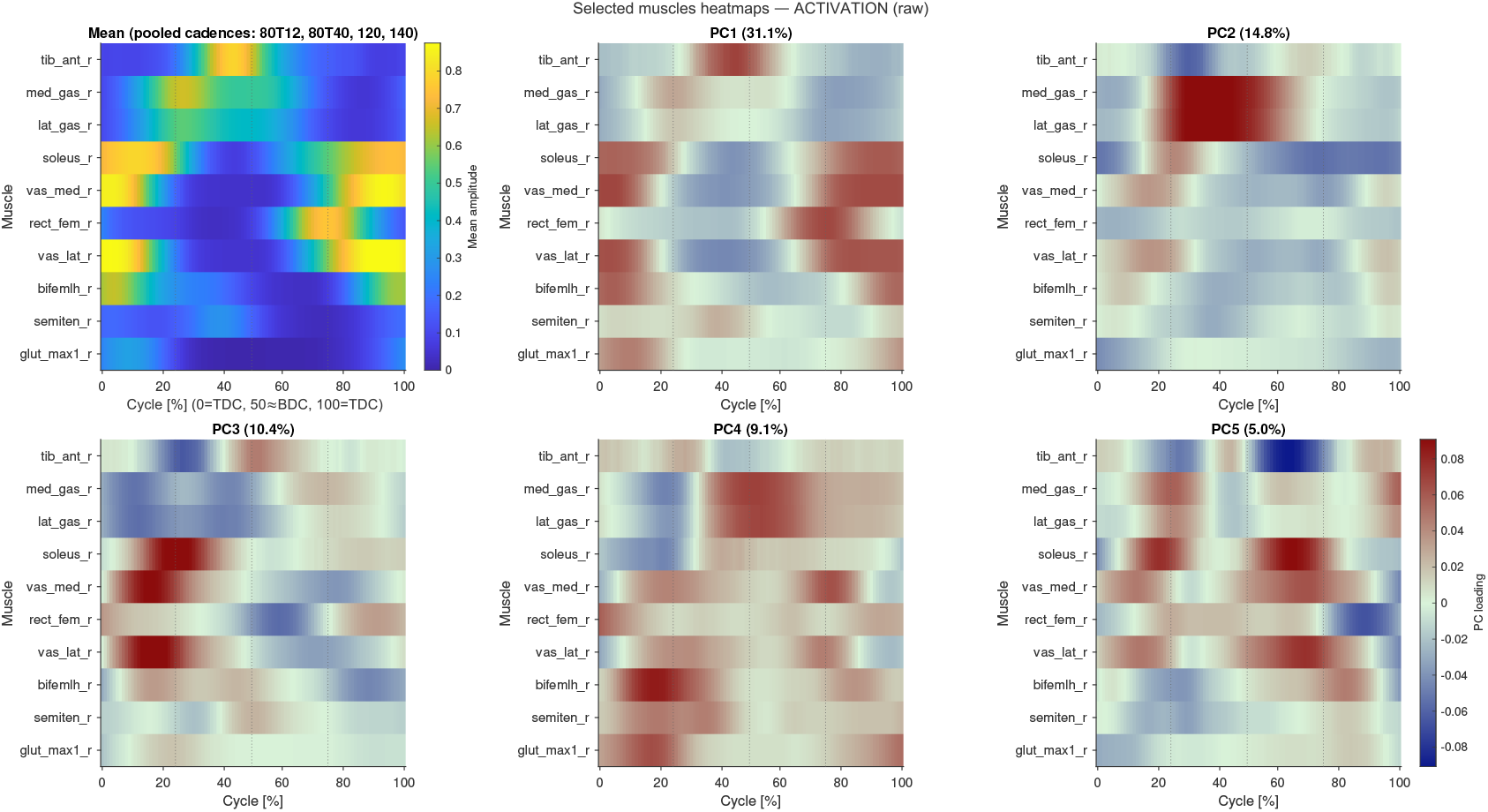
The mean activation pattern (A) and the weights for the first five Principal Components (PCs: 1-5). The variance explained is shown for each PC. Time is normalised to a percentage of the pedal cycle, starting and finishing with the crank at top-dead-centre.

The weights for PC1-5 accounted for changes to the coordination patterns between muscles, particularly within synergistic groups of muscle (Fig. 3). For instance, the presence of PC2 would promote more gastrocnemius but less soleus activity during the down-stroke of the pedal cycle, and PCs2 and 3 explain a greater activation in the vastii but a reduced activity in the rectus femoris of the quadriceps group just after top-dead-centre.

The first 25 PCs accounted for 90.74% of the variance of the coordination patterns, and mostly showed significant effects with mass and/or condition. These PCs were used to reconstruct the predicted activation patterns using the vector product of the PC loading scores (estimated marginal means or global mean) and the PC weights, summed back to the mean of the original activation matrix [64]. These reconstructed activation patterns were used to visualise the statistically significant main effects of the muscle mass and pedalling condition on the predicted activations and muscle coordination. ANOVA was conducted to determine the effect of the cycling condition and the muscle mass-scale for the loading scores of these first 25 PCs. There was a significant effect of cycling condition on the loading scores for 24/25 of these PCs, and a significant effect of muscle mass-scale on 11/25 of these PCs (Table S2). The effect of the muscle mass-scale on the loading scores of PC1-5 is shown in Figure. 4. The scale effects are most apparent for mass-scales of 70-125 and are clearly seen for the loading scores of PCs 2, 4, and 5. The effect of the cycling condition on the loading scores of PC1-5 is shown in Figure. 5. All five of these PCs show a strong effect of cadence.

**Figure 4.**
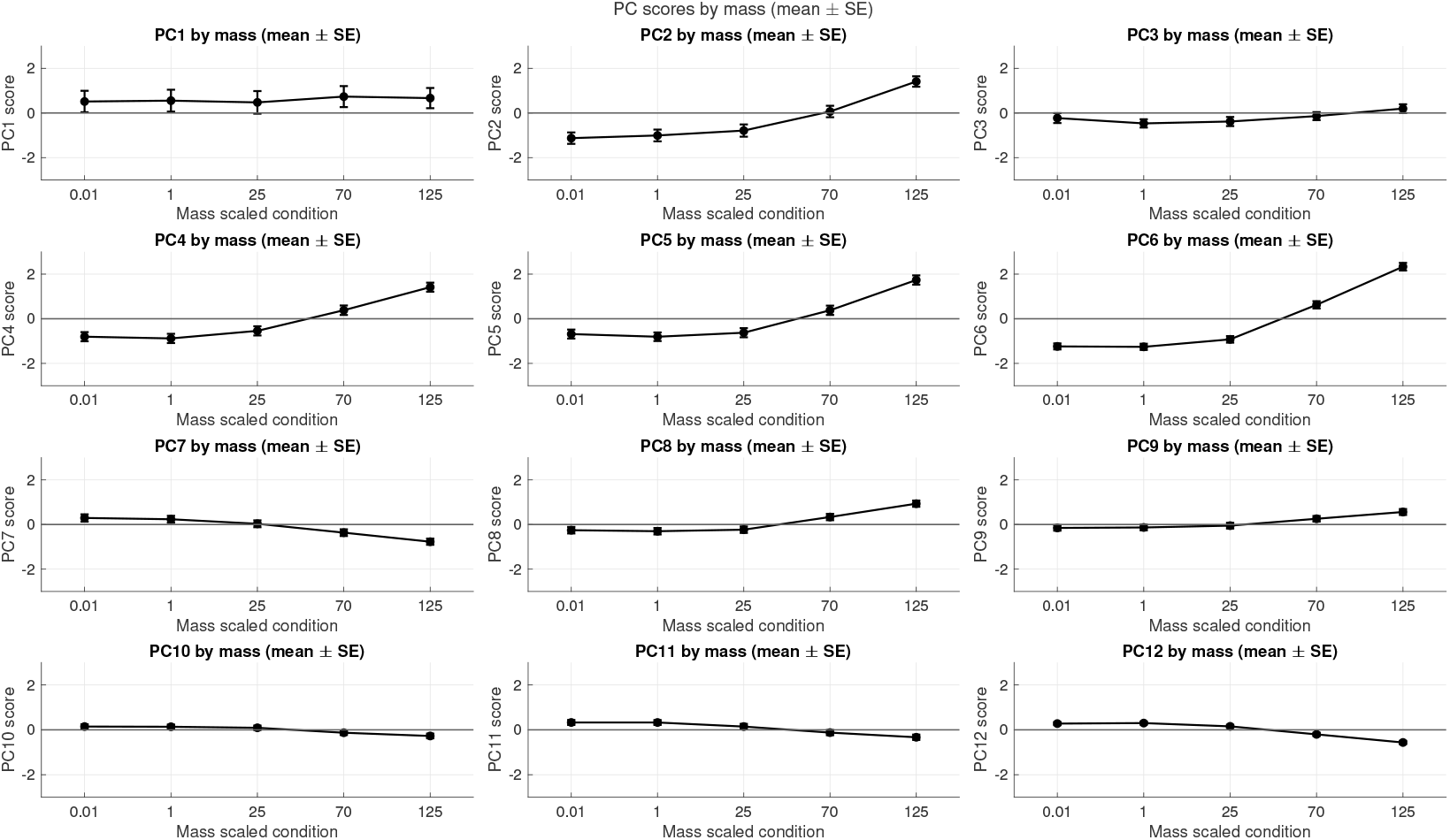
Loading scores for the effect of muscle-mass scale on the loading scores for PCs1-12. Symbols show the estimated marginal mean ± standard error of the mean from the ANOVA tests.

**Figure 5.**
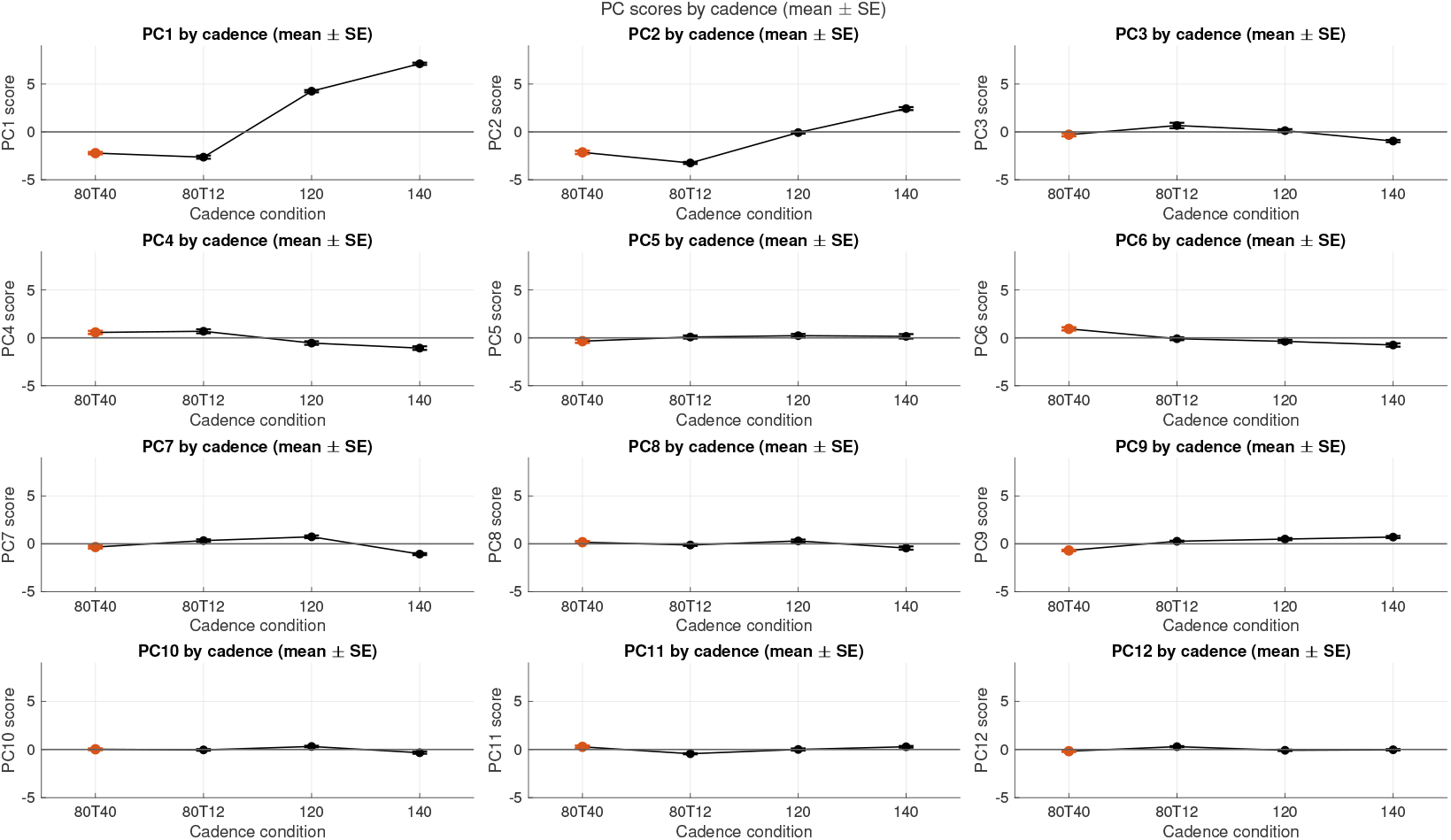
Loading scores for the effect of cycle condition of the loading scores for PCs1-12. Symbols show the estimated marginal mean ± standard error of the mean from the ANOVA tests.

The reconstructed activation patterns show that the muscle mass-scale causes a change in the magnitude of the activation-time profiles (Figure. 6), this is muscle-specific and is pronounced at scales 70-125.

**Figure 6.**
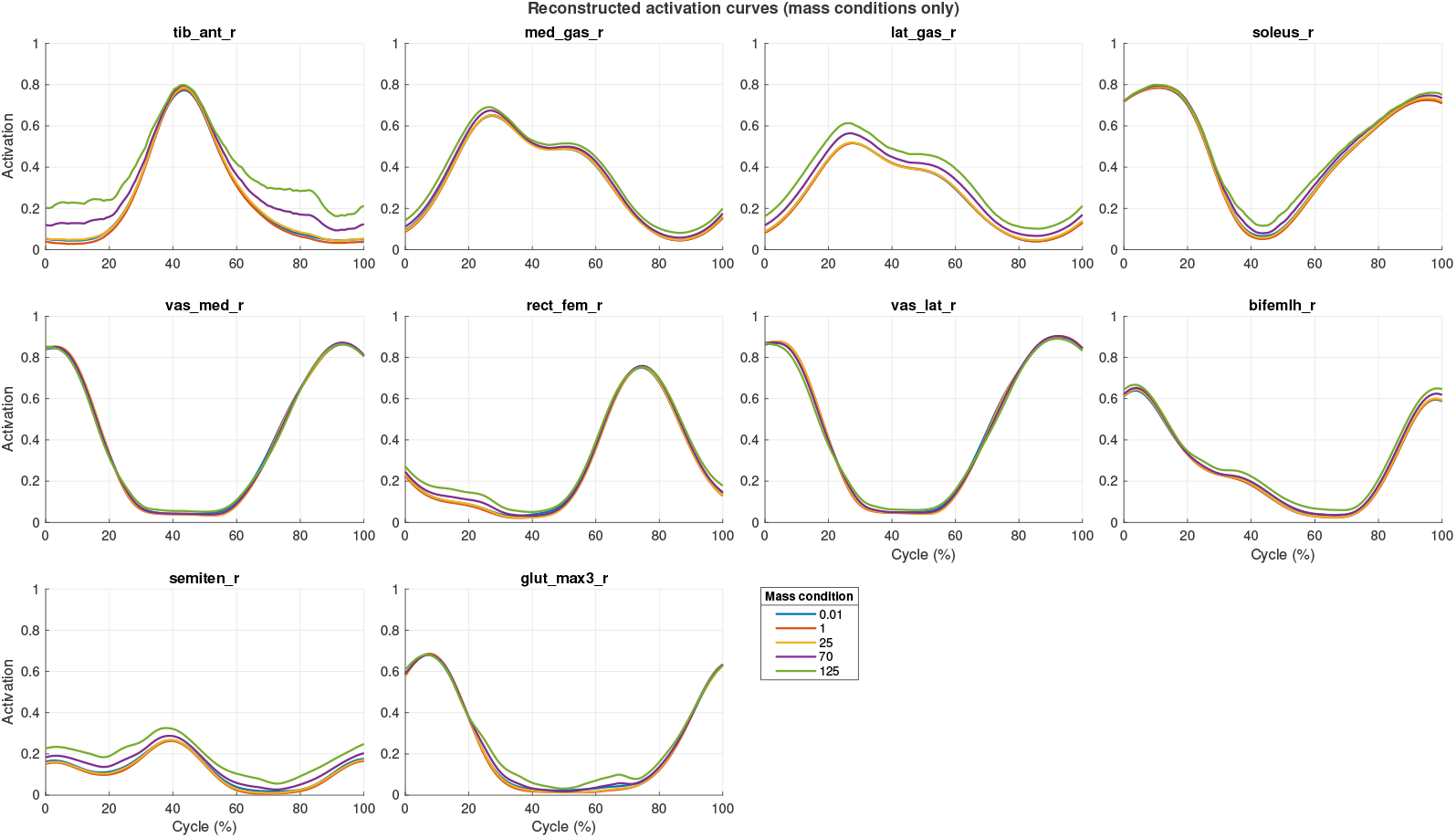
Reconstructed activation patterns for the select muscles using the mean activation and vector product of the loading scores and weights for PCs1-14. The statistical effect of the muscle-mass scale is shown. Loading scores were only selected to differ between conditions if there was a significant effect of scale for that PC from the ANOVA.

## Discussion

Musculoskeletal simulations showed that increased muscle mass in the simulations of pedalling resulted in differences in predicted muscle activations such that heavier muscles require greater activations and muscle mass significantly modulates the coordination patterns of the lower extremity muscles during cycling.

We have previously shown how the free-energy available for muscle contraction is distributed between internal and external work of the muscle [51]: muscle mass affects the contraction through both force balance and energy partitioning. Forces from the contractile elements contribute to accelerating the masses within the muscle. Motions of these masses thus have kinetic energy [77], with this energy being derived from the free energy available for contraction: thus, less energy will be available for the muscle to generate external work and power as the mass-scale increases. However, since the segmental kinematics, joint forces, and joint kinetics obtained in the inverse-dynamics simulations closely match the experimental data, the joint-level mechanical powers requirement remains essentially unchanged. Thus, we expected that the activation of the muscles would increase to achieve the required external work and power at the larger mass-scales because a higher proportion of the muscles’ free energy contributes to the internal kinetic energy. Hence, we expected that the muscle activation would increase to allow the required muscle power to be generated.

Forces from the contractile elements in the muscle models in these simulations had both force-length and force-velocity dependencies following their Hill-type model representations of the cross-bridge forces within the sarcomeres, with additional passive titin forces acting in parallel. Each muscle in the lower extremity has a unique pattern of muscle fibre strains and strain rates during locomotion. Thus, there would be unique set of accelerations, forces and powers for each muscle, and by extension the influence of muscle-mass on the muscle activations should also vary by muscle.

The size of the muscle-mass effects is small at the nominal human scale, as shown by the reconstructed activation patterns (Figure. 6). We present these reconstructed activations for a selection of the larger lower extremity muscles that make important contributions to the crank power during cycling. It should be noted that some of the smaller muscles (such as the flexor digitorum longus and flexor hallucis longus) showed larger mass effects on the predicted activations, but are not discussed here further due to their minor importance to cycling power (however, such smaller muscles may become more important with greater body size). A similarly small effect size was seen in our previous paper where we predicted the muscle forces in a muscle-by-muscle basis (for the same selected muscles as Figure. 3) in a forward dynamics model driven by experimentally measured EMG [22]. In both cases the muscles were represented by 1D serial lumped-mass models. These 1D models do not take into account the material properties or the 3D deformation of the muscle tissue. During contractions, muscle deformation occurs in 3D, and thus additional internal kinetic energy will be associated with this extra dimensionality and indeed, we have shown in both experimental and 3D modelling studies that muscle mass has an impact at scales even less than human size [13, 69]. Nonetheless, the physical laws of scaling are generalizable across the whole range of animal sizes, and the results from this study show that increasing muscle mass results in a general increase to the activations of the lower extremity muscles, in support of our first hypothesis.

In this study we quantified the Principal Components of the variance of the time-varying muscle activation patterns between the muscles. Activation traces (Figure 6) were reconstructed from the mean activation pattern and the vector-product of the Principal Component weights and loading scores. We enforced that the only difference in the reconstructions occurred when there were significant main effects of the muscle-mass scale or the cadence on the PC loading scores. Thus, differences in these reconstructed traces reflect significant effects of these factors. Each of the Principal Components contains both positive and negative weights of the activations (varying by muscle and time; Figure. 3). Thus, where the muscle mass-scale has a significant effect on these components then the relative muscle activity changes across muscles and time, and therefore these significant main effects of cadence and muscle mass-scale associate with modulations to the coordination pattern between the muscles. The loading scores for many of the PCs showed significant and monotonic changes with muscle mass-scale (Figure. 4), showing that modulations to the muscle coordination occur progressively with the change in muscle mass, in support of our second hypothesis.

The range of mass-scales selected for this study span the range of muscle masses for terrestrial vertebrates. However, we are not suggesting that the predicted coordination patterns that the simulations generated based on the cycling data would reflect the coordination across the range of animal sizes. Instead, we acknowledge that posture and gait vary considerably across the range of vertebrate sizes [36, 71, 72, 73], gait is very different between bipedal and quadrupedal animals [74, 75, 76] and differs between running and cycling. Additionally, cycling does not expose the body to foot-ground collisions with the associated tendency for soft-tissues to vibrate and wobble. However, the results point to the more general importance of muscle mass as a factor that affects the dynamics of muscle contraction, and thus the relative use of muscles for locomotion, with this being particularly the case for muscles that are larger than human-sized.

## Limitations

In interpreting our results, it is important to note that both model assumptions and the muscle-redundancy formulation can influence the predicted muscle-level responses. At the model level, Hill-type musculotendon dynamics depend on assumptions about activation–contraction dynamics, tendon compliance, and the force–length–velocity relations, which shape how joint-level demands are converted into fibre kinematics and tendon force transmission, particularly under rapid transients [1, 42, 81, 82]. Crucially, our “mass-scaling approach” (i.e., increasing muscle/segment mass and inertial properties without proportionally scaling geometric dimensions or force-generating capacity) is intentionally non-isometric: it isolates inertial effects but departs from physiological similarity laws, thereby altering the balance between inertial loads and available muscle force and potentially shifting muscles into different regions of their force–velocity curves.

More generally, scaling conventions based on body size and dynamic similarity [76] can change the normalization of forces, moments, and work and thus affect the magnitude of between-condition differences [43]. At the solver level, the choice of optimization criterion (e.g., minimizing activation/effort) when solving the muscle redundancy problem is known to bias load sharing between muscles; changes in objective weights, reserve-actuator penalties, or state bounds/regularization can also systematically redistribute recruitment across muscles even when the net joint moments are matched [44, 8]. Consequently, while qualitative trends may be robust, absolute magnitudes and the specific muscles showing increases versus decreases should be interpreted in light of these intertwined modeling, scaling, and optimization choices.

## Conclusion

This study presents the first use of explicit muscle inertia within 1D Hill-type muscle models for musculoskeletal simulations of movement. We found that the muscle mass modulates the predicted coordination pattern between the muscles; this effect was negligible at human mass-scales and became more pronounced at larger scales. The simulations predicted a general increase in muscle activation when the muscle mass-scale is equal to that of big terrestrial mammals. It is premature to conclude that the muscle mass is negligible at human size due to the simplification of the 1D models. Simulating mass effects with 3D muscle models may result in additional or larger effects.

## Acknowledgements

The authors express their gratitude to Dr. Friedl De Groote and Dr. Torstein E. Dæhlin, for their insightful helps about this study.

