## Supplementary Material for "The influence of muscle mass on the coordination required for efficient movement"

**Table S1.** Characteristics of the 12 participants.

| Label | Gender | Age (years) | Body mass (kg) | Height (m) |
|---|---|---|---|---|
| S02 | male | 28 | 68.0 | 1.67 |
| S03 | female | 30 | 65.5 | 1.67 |
| S04 | male | 33 | 68.0 | 1.75 |
| S05 | female | 43 | 64.4 | 1.74 |
| S06 | male | 32 | 63.0 | 1.83 |
| S07 | male | 37 | 71.8 | 1.73 |
| S11 | male | 31 | 82.8 | 1.83 |
| S12 | female | 24 | 65.8 | 1.71 |
| S13 | female | 32 | 63.5 | 1.68 |
| S14 | male | 24 | 78.9 | 1.78 |
| S15 | female | 26 | 58.0 | 1.67 |
| S17 | female | 26 | 80.7 | 1.73 |

**Table S2.** Statistical results for the GLM ANOVA for the PC loading scores for the first 25 PCs. Significants (p) is displayed for both the muscle-mass scale (Mass) and the pedal condition (Cond).

| PC | $F_{\text{Mass}}(4, 645)$ | $p_{\text{Mass}}$ | $F_{\text{Cond}}(3, 645)$ | $p_{\text{Cond}}$ |
|---|---|---|---|---|
| 1 | 0.25 | $p \geq 0.05$ | 350.50 | $p < 0.001$ |
| 2 | 20.58 | $p < 0.001$ | 71.96 | $p < 0.001$ |
| 3 | 0.69 | $p \geq 0.05$ | 15.18 | $p$<0.001 |
| 4 | 49.09 | $p < 0.001$ | 32.63 | $p < 0.001$ |
| 5 | 79.82 | $p < 0.001$ | 9.94 | $p < 0.001$ |
| 6 | 139.22 | $p < 0.001$ | 40.79 | $p < 0.001$ |
| 7 | 11.61 | $p < 0.001$ | 44.76 | $p < 0.001$ |
| 8 | 29.45 | $p < 0.001$ | 11.76 | $p < 0.001$ |
| 9 | 14.78 | $p < 0.001$ | 59.64 | $p < 0.001$ |
| 10 | 4.69 | $p < 0.001$ | 11.08 | $p < 0.001$ |
| 11 | 5.70 | $p < 0.001$ | 7.90 | $p < 0.001$ |
| 12 | 21.65 | $p < 0.001$ | 13.55 | $p < 0.001$ |
| 13 | 0.76 | $p \geq 0.05$ | 18.43 | $p < 0.001$ |
| 14 | 4.92 | $p < 0.001$ | 22.01 | $p < 0.001$ |
| 15 | 0.50 | $p \geq 0.05$ | 8.94 | $p < 0.001$ |
| 16 | 0.063 | $p \geq 0.05$ | 15.62 | $p < 0.001$ |
| 17 | 0.84 | $p \geq 0.05$ | 19.42 | $p < 0.001$ |
| 18 | 0.74 | $p \geq 0.05$ | 31.30 | $p < 0.001$ |
| 19 | 2.19 | $p \geq 0.05$ | 21.92 | $p < 0.001$ |
| 20 | 0.13 | $p \geq 0.05$ | 7.07 | $p < 0.001$ |
| 21 | 0.17 | $p \geq 0.05$ | 4.50 | $p < 0.05$ |
| 22 | 0.42 | $p \geq 0.05$ | 6.14 | $p < 0.001$ |
| 23 | 0.31 | $p \geq 0.05$ | 7.89 | $p < 0.001$ |
| 24 | 0.22 | $p \geq 0.05$ | 8.50 | $p < 0.001$ |
| 25 | 0.06 | $p \geq 0.05$ | 2.4 | $p \geq 0.05$ |

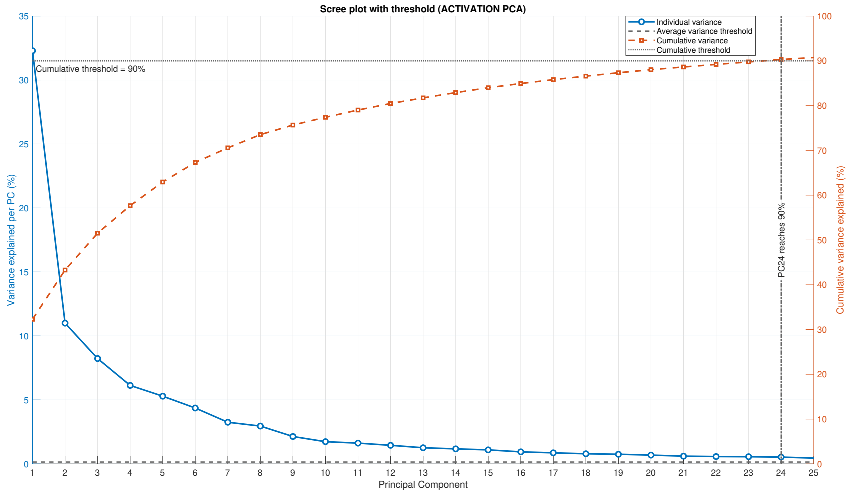


**Fig. S1.** Scree plot showing the percentage of the variance of the muscle coordination patterns explained by the first 25 PCs.

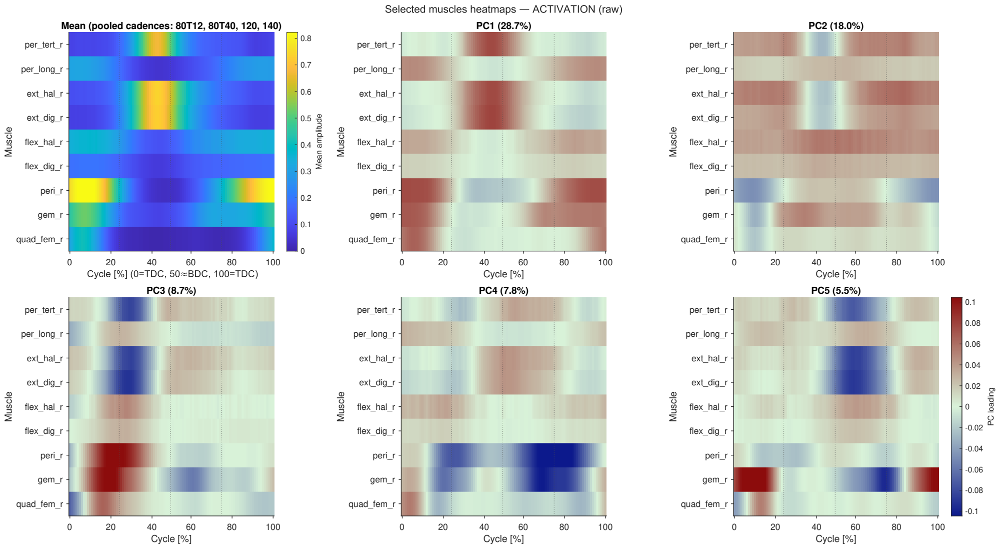


**Fig. S2.** The mean activation pattern and the weights for the first five Principal Components (PCs: 1-5). The variance explained is shown for each PC. Time is normalised to a percentage of the pedal cycle, starting and finishing with the crank at top-dead-centre.

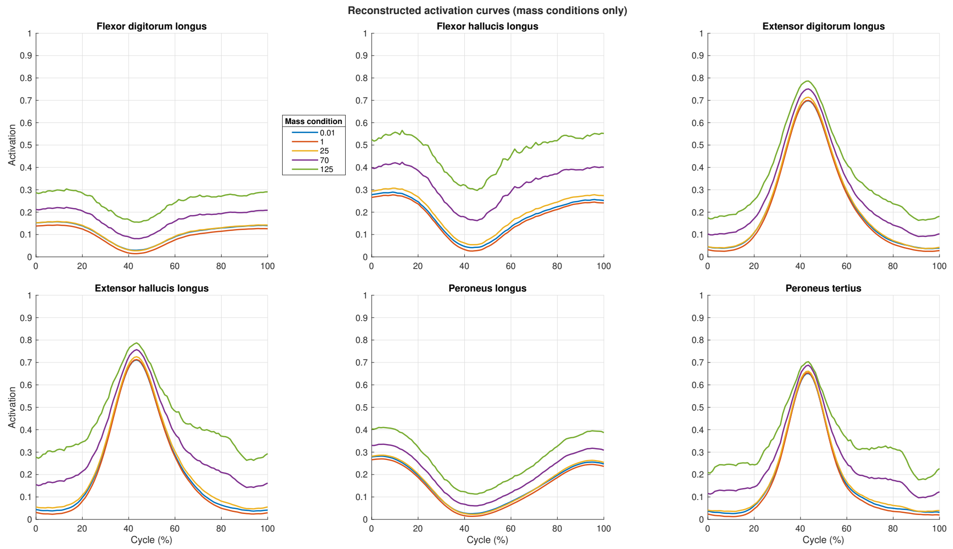


**Fig. S3.** Reconstructed activation patterns for the other muscles using the mean activation and vector product of the loading scores and weights for PCs1-14. The statistical effect of the muscle-mass scale is shown. Loading scores were only selected to differ between conditions if there was a significant effect of scale for that PC from the ANOVA.
